# GA-repeats on mammalian X chromosomes support Ohno’s hypothesis of dosage compensation by transcriptional upregulation

**DOI:** 10.1101/485300

**Authors:** Edridge D’Souza, Kathryn Weinand, Elizaveta Hosage, Steve Gisselbrecht, Vicky Markstein, Peter Markstein, Martha L. Bulyk, Michele Markstein

## Abstract

Over 50 years ago, Susumo Ohno proposed that dosage compensation in mammals would require upregulation of gene expression on the single active X chromosome, a mechanism which to date is best understood in the fruit fly *Drosophila melanogaster*. Here, we report that the GA-repeat sequences that recruit the conserved MSL dosage compensation complex to the *Drosophila* X chromosome are also enriched across mammalian X chromosomes, providing genomic support for the Ohno hypothesis. We show that mammalian GA-repeats derive in part from transposable elements, suggesting a mechanism whereby unrelated X chromosomes from dipterans to mammals accumulate binding sites for the MSL dosage compensation complex through convergent evolution, driven by their propensity to accumulate transposable elements.

## INTRODUCTION

Dosage compensation in placental mammals has long been known to involve epigenetic silencing of one of the two X chromosomes in females^1^, resulting in one silenced X chromosome and one active X chromosome. This mechanism ensures that the active X to autosome ratio is the same between females, which have two X chromosomes, and males, which have only one X chromosome in each somatic cell. In addition, a growing body of evidence suggests that dosage compensation in mammals involves a second mechanism that augments gene expression from the single active X chromosome so that it is equivalent to the output of two active X chromosomes^2-9^. This second mechanism was first proposed in 1967 by Susumu Ohno^10^, who argued that increased expression from the X chromosome would be required to avoid the consequences of aneuploidy arising from the evolutionary degeneration of its homolog into a gene-poor Y chromosome. However, while X-linked sequences that mediate the silencing arm of dosage compensation, such as Xist, have been identified, X-linked sequences that target genes on the active X chromosome for augmented expression have remained elusive.

Augmented expression of X-linked genes occurs by two mechanisms in mammalian cells: increased transcription and increased stabilization of RNA transcripts^7,9,11^. It is generally thought that these mechanisms evolved on a gene-by-gene basis on the X chromosome to compensate for the loss of homologous genes on the Y chromosome^7,12^. Consistent with this model, conservation of gene-specific microRNA target sequences correlates with the “dosage-sensitivity” of X-linked genes^13^. However, to date, no DNA sequences have been reported that support either a gene-by-gene or X-chromosome-wide mechanism resulting in augmented transcription of dosage-compensated genes in mammals.

Mechanistically, transcriptional upregulation of X-linked genes has been linked to the activity of the acetyltransferase MOF/KAT8, which acetylates histone 4 at lysine 16 (H4K16ac), resulting in open chromatin and increased transcription^14,15^. ChIP-seq experiments in mammalian cells show that MOF/KAT8, H4K16ac, and RNA Pol II are enriched about two-fold at upregulated X-linked genes relative to autosomal genes^7,9^. Moreover, RNAi knockdown of MOF/KAT8 in mammalian cells reduces the two-fold enrichment of RNA Pol II at several X-linked genes and their levels of transcription^9^. The question of how X-linked genes become targeted for transcriptional upregulation is therefore tied to how MOF/KAT8 becomes enriched at X-linked genes.

Since MOF/KAT8 plays a similar role in dosage compensation in the fruit fly *Drosophila melanogaster*^16,17^, we reasoned that its mechanism of enrichment might lend insight into how X-linked genes are targeted for upregulation in mammals. In both flies and mammalian cells, MOF/KAT8 is recruited to the X chromosome as part of the MSL dosage compensation complex, which contains the conserved proteins MSL1, MSL2, and MSL3^14-17^. In flies, these proteins have been shown to direct the MSL complex to the X chromosome in a two-step process: 1) MSL1 and MSL2 are required for recruitment of the complex to about 300 “chromatin entry sites” (CES), also called high affinity sites (HAS), along the X chromosome, and 2) MSL3, a chromodomain protein, is required for the spreading of the complex from the CES/HAS sites to neighboring genes^18-20^. In addition, two zinc finger proteins, CLAMP and GAF, recruit the MSL complex to DNA^21,22^. CLAMP acts locally at CES/HAS sites and distally with GAF to recruit the MSL complex and to shape the overall architecture of the X chromosome^22-24^. While it is unclear if there is a CLAMP homolog in mammals, molecular modeling has identified c-krox8/Th-POK/ZBTB7B as a mammalian GAF homolog^25^.

Analysis of the 300 recruitment CES/HAS sites in *Drosophila*, as well as *in vitro* binding assays with MSL2, CLAMP, and GAF, point to the importance of GA-dinucleotides in recruiting the MSL complex to the X chromosome. For example, analysis of the CES/HAS sites identified a 21-bp “MSL Recognition Element” (MRE) containing an 8-bp GA-repeat core that is necessary for MSL complex recruitment^18^. *In vitro* binding studies show that MSL2 binds a MRE-like sequence, called “PionX” that while different from the 5’ and 3’ ends of the MRE, retains the 8-bp GA-repeat core^26,27^. Additionally, *in vitro* binding assays with CLAMP show preferential binding to longer GA-repeats between 10 and 30 bp^23^.

## RESULTS

### GA-repeats are enriched along the human X chromosome and at X-linked dosage compensated genes

To determine if the density of GA-repeats is likewise enriched on mammalian X chromosomes and at mammalian X-linked genes, we developed “GenomeHash,” an algorithm to count user-defined motifs throughout specified genomes (see Methods). We validated our algorithm against the *Drosophila* genome, confirming previously published findings that the densities of GA-repeats are generally enriched ≥2-fold on the X chromosome relative to autosomes (Supplemental Table I). For example, GA-repeat lengths from 8 to 28 bp, experimentally validated in MSL and CLAMP binding assays *in vitro* and *in vivo*^18,23,27^, occur on the X chromosome at densities greater than once per million base pairs (≥1.0/Mb), with an average X chromosome to autosome (X:A) density enrichment of 2.5-fold (*p = 5.72*10^−118^ to 5.91*10^−07^, Poisson test)* (Figure 1a).

**Fig 1.**
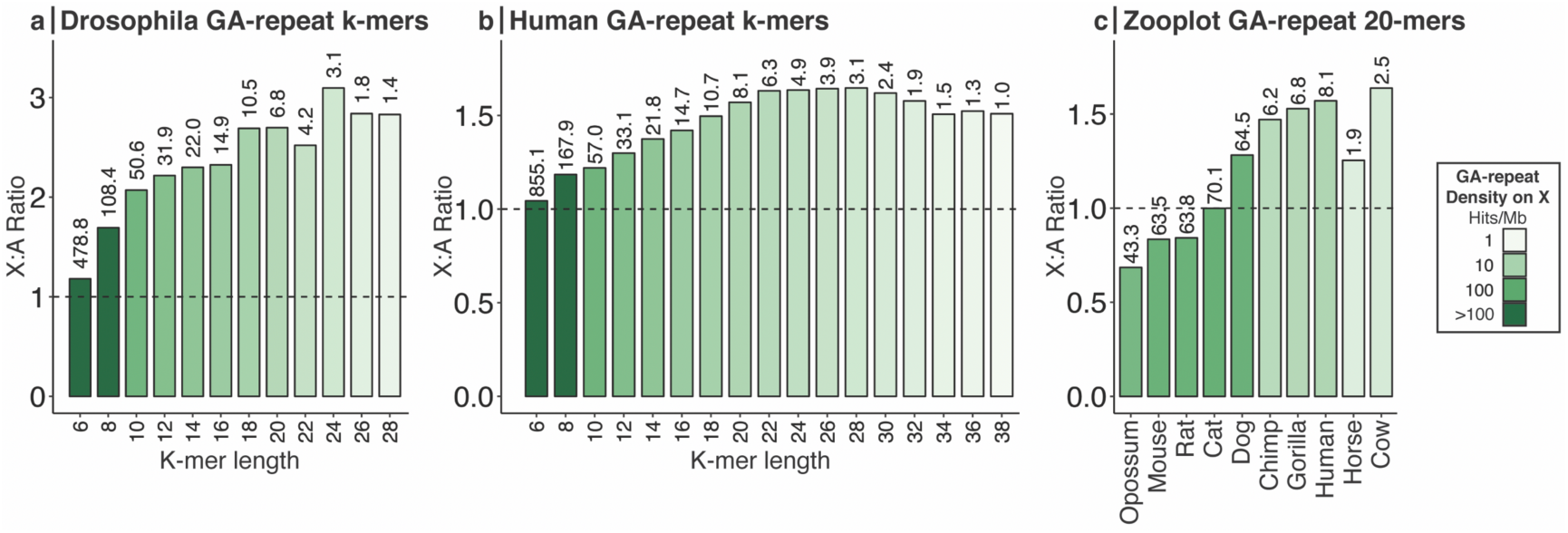
Density of GA-repeats on the X chromosomes of flies and mammals. The density of GA-repeats per megabase (Mb) was computed on each chromosome by dividing the number of GA-repeat matches (hits) by the length of each chromosome. The y-axis shows the ratio of GA-repeat densities on the X chromosome vs. autosomes for GA-repeats of specific lengths (X:A Ratio). Numbers above each bar represent the density of matches for each GA-repeat k-mer on the X chromosome, which as shown by the color-key, span 3 orders of magnitude. **a**. GA-repeat k-mers in the *Drosophila melanogaster* genome with an X chromosome density ≥ 1 hit/Mb show on average 2.5- fold X:A density enrichment. **b.** GA-repeat k-mers in the human genome with an X chromosome density ≥ 1 hit/Mb show an average 1.5-fold X:A density enrichment. **c.** The density of GA-repeats of length 20 bp across different mammalian genomes. Genomes with low X:A enrichment ratios tend to have high baseline densities of GA- repeats.

We found that the density of GA-repeats on the human X chromosome is likewise enriched relative to autosomes *(p = 2.26*10^−72^ to 0.03 for 1.3-fold and above enrichment, Poisson test)*. Most prominently, GA-repeats of lengths 18-38 bp occur on the X chromosome at densities ≥ 1.0/Mb, where they are enriched 1.5-fold relative to autosomes (Figure 1b, Supplemental Table 1). Although the X:A enrichment of GA-repeat densities is more modest than that observed in *Drosophila*, it is statistically significant based on Poisson tests and empirical bootstrapping methods (see Materials and Methods) with Poisson test p-values ranging from 1.84*10^−57^ for 18-mers to 1.82*10^−07^ for 38-mers (Supplemental Table 1). Moreover, we found that dosage-compensated genes^28^ are more likely than autosomal genes to contain intronic GA-repeats (Supplemental Tables 2 and 3). For example, GA-repeats of length 30 bp occur in 12% of dosage dosage-compensated genes but only 5.3% of autosomal genes, resulting in a 2.2-fold enrichment of dosage-compensated genes (*p = 2.72*10^−8^, Poisson test*). We found similar results with clusters of GA-rich consensus motifs matching the *Drosophila* MSL recognition element (MRE) as well as binding sites for the *Drosophila* CLAMP protein and the mammalian GAF homolog (Supplemental Tables 2). Collectively, these findings show that the density and distribution of GA-repeats on the human X chromosome, as in *Drosophila*, are compatible with mediating chromosome-wide and gene-by-gene mechanisms of dosage compensation.

### GA-repeats are enriched along mammalian X chromosomes in two patterns

Further support that GA-repeats may mediate dosage compensation in mammals, stems from our finding that GA-repeats are enriched on the X-chromosome not only in the human genome but also the chimpanzee, gorilla, dog, cat, cow, horse, mouse, rat, and opossum genomes (Supplemental Table 1). Focusing on 20-mer GA repeats, two patterns of enrichment are apparent (Figure 1c). One pattern follows the ∼1.5-fold X:A density enrichment of GA-repeats that is observed in humans, and is evident in dogs, primates, horses, and cows. The second pattern is characterized by high densities of GA-repeats genome-wide. These higher densities of GA-repeats are about an order of magnitude greater than found on the *Drosophila* X chromosome and are found throughout the genomes of opossum, mouse, rat, cat and dog. With the exception of the dog genome, ultra-dense long GA-repeats in these genomes tend not to be enriched on the X chromosome.

Importantly, the densities of long GA-repeats across mammalian X chromosomes are statistically significant (Poisson tests and bootstrapping methods) (Supplemental Tables 2 and 3) and on par with other biologically active motifs that shape whole chromosomes. In fact, the occurrences of 20-bp GA-repeats on all 10 mammalian X chromosomes are more dense than the well-characterized genome-wide insulator protein, CTCF, which occurs about once every million base pairs across human chromosomes^29^. Additionally, 8 of the 10 mammalian genomes that we examined exhibit X chromosome GA-repeat densities on par with or greater than the density of GA-repeats associated with dosage compensation in *Drosophila*^23^.

### GA-repeats in mammalian genomes derive in part from transposable elements

Our finding that GA-repeats are abundant on X chromosomes across mammals supports the hypothesis that the MSL dosage compensation complex, known to interact with GA-repeats on the X-chromosomes in dipterans to augment gene expression, engages in a similar mechanism in mammals. However, unlike the core proteins of the MSL complex, which are derived from the last common ancestor between mammals and dipterans, GA-repeats cannot have arisen from a common ancestor, as the X chromosomes in dipterans and mammals have completely different evolutionary histories. In fact, even within dipterans, the X chromosomes have different evolutionary histories^30^, and have been shown in one case to acquire 8-bp GA-repeats through invasion and domestication of a transposable element (TE)^31^.

We likewise identified several examples of TEs associated with GA-repeats in the human genome. For example, LINE subfamilies L1 and L2, and the SINE subfamilies AluJ, AluS, AluY, have contributed loci with tandem duplications^32^ containing GA-repeats ranging from 8 to 28 bp (Supplemental Table 4). These findings suggest that further investigation of LINEs^33^, some of which exhibit a 2-fold X:A enrichment and SINEs^34^, present in high copy numbers in the genomes of opossum, mouse, rat, and dog, may explain the two patterns of GA-repeat enrichment we observed in mammalian genomes (Figure 1c). Additionally, other TE families may have also contributed to GA-repeat enrichment on the X chromosome; for example, we discovered that the mammalian gypsy retrotransposon (mam-Gypsy) is a good candidate. Based on the Hidden Markov Models (HMMs) available through the DFAM online database^35^, we found that mam-Gypsy retrotransposons contain G/A rich tracts in their Long Terminal Repeats (Figure 2a) and are three-fold enriched on the human X chromosome relative to autosomes (*p = 6.14*10^−23^, Poisson test*) (Figure 2b). These findings suggest that systematic exploration of TEs is likely to shed light on the evolution of GA-repeats in mammalian genomes.

**Fig 2.**
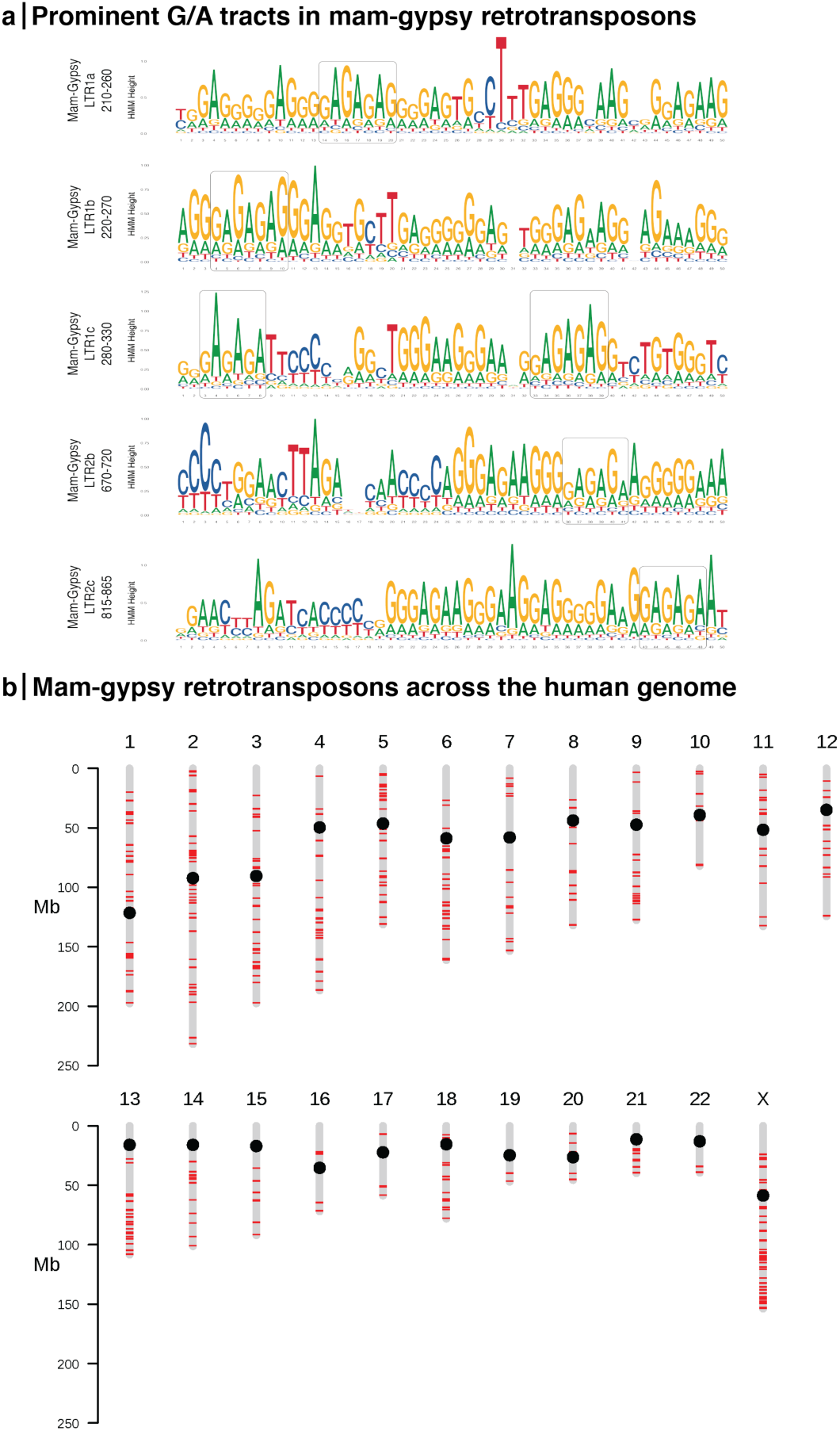
Distribution of the mammalian Gypsy retrotransposon in the human genome. The mam-Gypsy retrotransposon contains long G/A tracts and is enriched on the X chromosome. **a.** Logos showing G/A-rich tracts, spanning up to 50 bp, in mam-gypsy LTR regions, based on DFAM^35^ Hidden Markov Models. Boxed sequences show GA-repeat regions. **b.** Ideogram of genomic locations matching the DFAM^35^ database mammalian gypsy consensus models. The density of matches on the X chromosome is 3-fold higher than the mean autosomal density (p=6.14e-23, 1-tailed upper Poisson test).

## DISCUSSION

Collectively, our results show that just as the MSL dosage compensation complex is conserved from flies to mammals, its corresponding GA-rich core recognition motif is enriched on the X chromosomes of flies and mammals by convergent evolution. Our finding that GA-repeats in the human genome are derived in part from TEs suggests that any chromosome with a propensity to accumulate TEs and repetitive elements, such as ancient and nascent X chromosomes^31,33,36^, is poised to be targeted by transcriptional machinery, such as the MSL complex. Furthermore, we predict that the MSL complex, by virtue of its recruitment to chromosomes by GA-repeats, is poised to target future X chromosomes that may arise in evolution, as our results show that accumulation of GA-repeats is a common feature of X chromosomes.

## Methods

Software, Statistical Methods, and Databases employed are provided in the Supplemental Materials and Methods.

## Acknowledgements

We are indebted to Erica Larschan and Courtney Babbitt for insightful conversations and Jake Mayfield, Laura Quilter, Rachel Brody and Kristopher Kolbert for critically reviewing the manuscript. This project was supported by NIH grant R01 HG009723 to M.L.B. K.W. was supported in part by NIH Bioinformatics and Integrative Genomics training grant T32 HG002295. E.D. was supported in part by a Goldwater Scholarship and UMass Honor College Research Grant.

## Author contributions

M.M. and E.D. conceived the project; E.D., K.W., E.H., S.G., M.B., and M.M. designed the analyses; E.D., K.W., E.H., S.G. performed the analyses; V.M. and P.M. wrote GenomeHash; E.D. and K.W. generated the figures; K.W., S.G., and M.B. devised and performed bootstrapping statistical methods; E.D. and K.W. wrote the materials and methods section; M.M. wrote the manuscript; all authors edited and provided comments on the manuscript; M.B. and M.M. supervised the project.

