## Supplemental Materials and Methods for "GA-repeats on mammalian X chromosomes support Ohno’s hypothesis of dosage compensation by transcriptional upregulation"

[4]in silico Labs, 160 Redland Road, Woodside, CA 94062

[5]Department of Pathology; Brigham and Women’s Hospital and Harvard Medical School, Boston, MA 02115

### Supplemental Materials and Methods

#### Genomes

All genomes were obtained from UCSC Genome Browser[1] (https://genome.ucsc.edu/). Genome versions AgamP4, bosTau6, canFam3, dm6, equCab2, felCat8, galGal6, gorGor4, mm10, monDom5, panTro5, and rn6 were used for cross-species comparisons. Human genome version hg38 was used in all analyses except where noted, when instead hg19 was used. Gene annotations were downloaded in BED format from the UCSC Table Browser under the track “NCBI RefSeq.” A curated list of

348 X-linked dosage compensated genes was obtained from Tukiainen et al. (2017)[2] Supplement Table 13.

**Packages**

Third-party software packages were used for genome manipulation and visualization. We used the Python package mygene to convert gene ID formats (https://pypi.org/project/mygene/), as well as BioPython (https://biopython.org/) to access the sequences defined in the genomic BED files. The Python packages numpy (http://www.numpy.org/), matplotlib (https://matplotlib.org/), pandas (https://pandas.pydata.org/), and scipy (https://www.scipy.org/) were used for numerical computation and visualization. The R package chromPlot (https://rdrr.io/bioc/chromPlot/) was used to visualize the positions of transposable elements in the genome, and final plots were generated using the R package ggplot2 (https://ggplot2.tidyverse.org/). To determine which motifs and transposable elements are associated with genes, we used the command-line utility bedtools (https://github.com/arq5x/bedtools2) on BED files denoting coordinates of genes, motif search results, and transposable elements.

**Software developed**

We developed custom scripts in Bash and Python 3.5.5 to automate the analysis pipeline, as well as a custom search program GenomeHash (https://github.com/marksteinlab/GenomeHash). For a given motif, GenomeHash outputs genomic coordinates for all matches. A cluster is parametrized by its search sequence, the minimum number of matches of the search sequence within the cluster, and the maximum window size of the cluster. The coordinates outputted from

GenomeHash were correlated against chromosome, gene, and transposable element coordinates to assess the association between motifs and known feature annotations.

**Motif density analysis**

We calculated the overrepresentation of GA dinucleotide motifs on the X chromosome by running GenomeHash on GA repeats of length 2 through 60. The reported results for each motif were restricted to “clusters” that contain at least one match within a window length equal to the length of the search sequence. This is equivalent to searching for all occurrences of a given motif within the genome, without counting overlapping motifs. For example, in a GA-repeat of length 40 bp, our algorithm recognizes two 20 bp-repeats, one from positions 1-20 and the other from positions 21-40; the algorithm ignores the 8 overlapping repeats from positions 2-21, 3-22, 4-23, etc.

We reported the density of each motif as the frequency of matches within a given genomic region. To calculate the overrepresentation of different motifs on the X chromosome, we defined the “X density” as the number of matches on X divided by the length of the X chromosome, in Mb. We calculated “A density” similarly as the total number of autosomal matches divided by the total autosomal length, in Mb. The X:A density ratio divides these two terms in order to find how overrepresented a given motif is on X. In each case, we assumed a null hypothesis that the X density was less than or equal to the A density, and used a 1-tailed upper Poisson test to assess significance. This process was the same when comparing different motifs within a single species, and when comparing motifs across different animal species.

**Neighborhood analysis**

To determine how motifs of interest associate with X dosage compensated genes, we performed both static "neighborhood analysis" as well as empirical bootstrapping, as described below. The static neighborhood analysis compares a 'foreground' of X dosage compensated genes and a 'background' of autosomal genes. We extended both the foreground and background by the same flank length in order to obtain a "gene neighborhood"; for each motif in a list of query motifs, we determined the percentage of dosage compensated neighborhoods and the percentage of autosomal neighborhoods that contain a match for the target motif. We repeated this for flank sizes of 0, 1, 5, 10, and 30 Kb. For each search motif at each flank size, we tested whether the proportion of dosage compensated neighborhoods was higher than the proportion of autosomal neighborhoods using a 1-tailed upper binomial test. When testing multiple search sequences at once, we adjusted our significance threshold using Bonferroni correction and a family-wise error rate of 0.05.

**Transposable element analysis**

We obtained Hidden Markov Models of mam-Gypsy from the Dfam database[3] (http://www.dfam.org/) and we used ggseqlog[4] to create sequence logos of the G/A tracts. We used the Dfam database[3] to retrieve a list of all matching genomic coordinates for a given TE in both mice and humans. For each matching region, we accessed the raw sequence from the hg38 and mm10 genomic FASTA files and then calculated summary statistics for GA content. To assess whether each transposable element is biased toward the X chromosome, we used a process similar to the motif density analysis. X density and A density were calculated as the number of matches divided by the length of their respective search spaces, and the X:A ratio of these two densities indicates the degree to which a TE is biased towards the X chromosome. To test the significance of this bias, we performed a 1-tailed upper Poisson test, as with the motif density analysis.

A list of transposable elements with tandem repeats was obtained from Ahmed and Liang (2014)[5], Supplement Table 4.

**Bootstrap analysis**

To calculate the significance of the association between GA repeats and human genes, X dosage compensated genes, DNase-seq peaks, and ChIP-seq peaks (i.e. foreground regions), we used permutation testing. We did this for the gene body or peak regions themselves as well as flanking regions: gene TSS ± 3kb, 10kb, 30kb; gene TTS ± 3kb, 10kb, 30kb; peak center/summit ± 1kb, 10kb. Each dataset was filtered to exclude regions from chromosome Y, the mitochondrial chromosome, and any contigs. Regions were merged within each dataset to avoid double counting. All regions in this bootstrap analysis were mapped to the hg19 reference genome.

The first type of analysis compared chromosome X regions to autosomal chromosome regions within datasets. We generated 1000 randomly sampled autosomal chromosome background sets that were matched for chromosome X foreground set length distribution and set size.

The second type of analysis compared the whole dataset to randomly chosen length- and chromosome X-matched background sets. By chromosome X-matched, we mean that if the foreground region was on chromosome X, then the background region must also be on chromosome X; the same was done for the autosomal chromosomes. We were unable to generate a background set for the whole gene body datasets due to sequence scarcity. For the DNase-seq and ChIP-seq peak datasets, we generated 1000 background sets as described above, building off a GENRE database structure[6]. For the flanking regions, we encountered sequence scarcity for generated backgrounds after merging regions (region length became prohibitive). We therefore randomly sampled 1000 foreground subsets, picking only one foreground region from each group of overlapping regions, leaving a non-overlapping foreground set the size of which determined by how many backgrounds could be generated. A single background was generated as described above. The bootstrap script was run with a flag to compare 1000 foregrounds to 1 background.

A bootstrap was run to compare the density of GA repeat kmers in foreground regions to the background regions generated by each method. Three different lengths of GA repeat kmers were used: 20mer, 30mer, and 40mer. FASTA files were generated for each foreground and background set in two ways: one with CDS regions masked and one with nothing masked. An empirical p-value is reported as determined by the number of times a background set has greater or equal the number of GA kmers than the foreground set divided by the number of sets iterated over. For the whole dataset flanking region analysis, foreground sets were iterated over; all other analyses were background sets. An FDR-correction was performed to address the multiple hypothesis testing.

**Statistical testing**

As addressed above, we used both static and sampling-based methods to obtain the significance of different hypothesis tests. To determine whether motif matches from GenomeHash were overrepresented on X, we used a 1-sided upper Poisson test, which reduces to a simple binomial test comparing the density of hits on the X chromosome versus the autosomes. We used the same process to determine whether TE matches were biased toward the X chromosome. For the neighborhood analysis, we used a 1-sided upper binomial test to test whether the proportion of gene neighborhoods containing sequence matches was higher on the X chromosome than on the autosomes. We used Bonferroni correction to adjust the resultant significance threshold, holding a family-wise error rate of 0.05.

For the bootstrapping procedures, p-value for each search was calculated as the proportion of the 1000 shuffled backgrounds that exhibited sequence count greater than or equal to that of the foreground. Multiple hypothesis correction was done by using the p.adjust() function in R to yield Benjamini-Hochberg FDR adjusted p-values.

To establish that GA-repeats are overrepresented on the X chromosome, we performed another bootstrap analysis, searching the human genome for a GA 20-mer as well as 999 other shuffled iterations of the sequence. We obtained a list of sequences that were overrepresented on the X chromosome using the motif density analysis procedure. Of these sequences, we established the GA 20-mer as an outlier by comparing its density to the density of all the 73 total shuffled sequences that were also present on the X chromosome. We further established the significance of the GA 20-mer as an outlier by Studentizing its density value and comparing it to a T distribution with 72 degrees of freedom.
