## Supplement Table 2 for "GA-repeats on mammalian X chromosomes support Ohno’s hypothesis of dosage compensation by transcriptional upregulation"

| Motif | Cluster | % X DC genes | % A genes | X:A ratio | p-value |
|---|---|---|---|---|---|
| **GA 16mer** | Single motif | 32.6% | 18.8% | 1.74 | 7.80E-12 |
| **GA 18mer** | Single motif | 27.2% | 14.9% | 1.83 | 2.47E-11 |
| **GA 20mer** | Single motif | 23.5% | 12.3% | 1.91 | 5.70E-11 |
| **GA 30mer** | Single motif | 11.9% | 5.4% | 2.20 | 2.72E-08 |
| **GA 40mer** | Single motif | 5.4% | 2.3% | 2.30 | 6.93E-05 |
| **GA 50mer** | Single motif | 2.7% | 0.8% | 3.31 | 4.12E-05 |
| **dCLAMP consensus** | Single motif | 49.6% | 34.1% | 1.45 | 2.37E-10 |
| **dCLAMP consensus** | 3 motifs in 100 bp | 9.2% | 3.9% | 2.36 | 1.12E-07 |
| **dMRE consensus** | Single motif | 66.6% | 51.2% | 1.30 | 1.94E-09 |
| **dMRE consensus** | 3 motifs in 100 bp | 13.7% | 7.4% | 1.85 | 2.45E-06 |
| **mGAF consensus** | Single motif | 82.2% | 68.7% | 1.20 | 1.14E-08 |
| **mGAF consensus** | 3 motifs in 100 bp | 12.9% | 6.5% | 2.00 | 2.90E-07 |

**Supplementary Table 2: GA-repeat motifs in the gene bodies of X-linked dosage compensated (DC) genes.** Matches are counted if they lie within the body of a gene. "% DC genes" refers to the percent of 348 curated dosage-compensated (DC) genes containing at least one corresponding motif; "% Autosomal genes" refers to the percent of autosomal genes containing at least one corresponding motif; "X:A" is the ratio of the preceding two columns' "p-value" refers to upper-tailed binomial p-values. Using Bonferroni correction with $\alpha = 0.05$ and 11 comparisons, the critical p-value is 4.54E-3.
*Drosophila* dCLAMP consensus : RAGAGARMGAGANRR
*Drosophila* dMRE consensus: RWNNGARMGAGANRR
Mammalian mGAF consensus: VVKHD-DGARVVHBRVDBRRMR
